# Faucet: streaming de novo assembly graph construction

**DOI:** 10.1101/125658

**Authors:** Roye Rozov, Gil Goldshlager, Eran Halperin, Ron Shamir

## Abstract

**Motivation:** We present Faucet, a 2-pass streaming algorithm for assembly graph construction. Faucet builds an assembly graph incrementally as each read is processed. Thus, reads need not be stored locally, as they can be processed while downloading data and then discarded. We demonstrate this functionality by performing streaming graph assembly of publicly available data, and observe that the ratio of disk use to raw data size decreases as coverage is increased.

**Results:** Faucet pairs the de Bruijn graph obtained from the reads with additional meta-data derived from them. We show these metadata - coverage counts collected at junction k-mers and connections bridging between junction pairs - contain most salient information needed for assembly, and demonstrate they enable cleaning of metagenome assembly graphs, greatly improving contiguity while maintaining accuracy. We compared Faucet’s resource use and assembly quality to state of the art metagenome assemblers, as well as leading resource-efficient genome assemblers. Faucet used orders of magnitude less time and disk space than the specialized metagenome assemblers MetaSPAdes and Megahit, while also improving on their memory use; this broadly matched performance of other assemblers optimizing resource efficiency - namely, Minia and LightAssembler. However, on metagenomes tested, Faucet’s outputs had 14-110% higher mean NGA50 lengths compared to Minia, and 2-11-*fold* higher mean NGA50 lengths compared to LightAssembler, the only other streaming assembler available.

**Availability:** Faucet is available at https://github.com/Shamir-Lab/Faucet

**Supplementary information:** Supplementary data are available at *Bioinformatics* online.

## 1 Introduction

Assembly graphs encode relationships among sequences from a common source: they capture sequences as well as the overlaps observed among them. When assembly graphs are indexed, their sequence contents can be queried without iterating over every sequence in the input. This functionality makes graph and index construction a prerequisite for many applications. Among these are different types of assembly - e.g., de novo assembly of whole genomes, transcripts, plasmids, etc. [1, 2] - and downstream applications - e.g., mapping reads to the graphs, variant calling, pangenome analysis, etc. [3, 4]

In recent years, much effort has been expended to reduce the amount of memory used for constructing assembly graphs and indexing them. Major advances often relied on index structures that saved memory by enabling subsets of possible queries: e.g., one could query what extensions a given substring *s* has, but not how many times *s* was seen in the input data. A great deal of success ensued in reducing the amount of memory needed to efficiently construct the central data structures used by most de novo assembly algorithms, namely, the de Bruijn and string graphs [5, 6, 7, 8]. Furthermore, efficient conversion of de Bruijn graphs to their *compacted* form (essentially string graphs with fixed overlap size) has been demonstrated [9, 10, 11].

In parallel to these efforts, streaming approaches were demonstrated as alternative resource-efficient means of performing analyses that had typically relied on static indices. Although appealing in terms of speed and low memory use, these approaches were initially demonstrated primarily for counting-centered applications such as estimating k-mer frequencies, error-correction of reads, and quantification of transcripts [12, 13, 14, 15, 16].

Recently, a first step towards bridging the gap between streaming approaches and those based on static index construction was taken, hinting at the potential benefits of combining the two. Matwali et al.[17] demonstrated a streaming approach to assembly by making two passes on a set of reads. The first pass subsamples k-mers in the de Bruijn graph and inserts them into a Bloom filter, and the second uses this Bloom filter to identify ’solid’ (likely correct) k-mers, which are then inserted into a second Bloom filter. This streaming approach resulted in very high resource efficiency in terms of memory and disk use. However, LightAssembler finds solid k-mers while disregarding paired-end and coverage information, and thus is limited in its ability to resolve repeats and to differentiate between different possible extensions in order to improve contiguity.

In this work, we extend this approach with the aim of providing a more complete alternative to downloading and storing reads for the sake of de novo assembly. We show this is achievable via online graph and index construction. We describe the Faucet algorithm, composed of an online phase and an offline phase. During the online phase, two passes are made on the reads without storing them locally to first load their k-mers into a Bloom filter, and then identify and record structural characteristics of the graph and associated metadata essential for achieving high contiguity in assembly. The offline phase uses all of this information together to iteratively clean and refine the graph structure.

We show that Faucet requires less disk space than the input data, in contrast with extant assemblers that require storing reads and often produce intermediate files that are larger than the input. We also show that the ratio of disk space Faucet uses to the input data improves with higher coverage levels by streaming successively larger subsets of a high coverage human genome sample. Furthermore, we introduce a new cleaning step called *disentanglement* enabled by storage of paired junction extensions in two Bloom filters - one meant for pairings inside a read, and one meant for junctions on separate paired end mates. We show the benefit of disentanglement via extensive experiments. Finally, we compared Faucet’s resource use and assembly quality to state of the art metagenome assemblers, as well as leading resource-efficient genome assemblers. Faucet used orders of magnitude less time and disk space than the specialized metagenome assemblers MetaSPAdes and Megahit, while also improving on their memory use; this broadly matched performance of other assemblers optimizing resource efficiency - namely, Minia and LightAssembler. However, on metagenomes tested, Faucet’s outputs had 14-110% higher mean NGA50 lengths compared to Minia, and 2-11-*fold* higher mean NGA50 lengths compared to LightAssembler, the only other streaming assembler available.

## 2. Preliminaries

For a string *s*, we denote by *s[i]* the character at position *i, s*[*i*: *j*] the substring of *s* from position *i* to *j* (inclusive of both ends), and *|s|* the length of *s*. Let *pref* (*s, j*) be the prefix comprised of the first *j* characters of *s* and *suff* (*s, j*) be the suffix comprised of the last *j* characters of *s*. We denote concatenation of strings *s* and *t* by *s ∘ t*, and the reverse complement of a string *s* by *s′*.

A *k-mer* is a string of length *k* drawn from the DNA alphabet Σ = *{A, C, G, T }*. The de Bruijn graph *G*(*S, k*) = (*V, E*) of a set of sequences *S* has nodes defined by consecutive k-mers in the sequences, 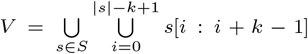; *E* is the set of arcs defined by (*k−* 1)– *mer* overlaps between nodes in *V*. Namely, identifying vertices with their k-mers, (*u, v*) *∈ E ⇔ suff* (*u, k −* 1) = *pref* (*v, k −* 1). Each node *v* is identified with its reverse complement *v′*, making the graph *G* bidirected, in that edges may represent overlaps between either orientation of each node [18]. When necessary, our explicit representation of nodes will use *canonical* node naming, i.e., the name of node (*v, v′*) will be the lexicographically lesser of *v* and *v′*. *Junction nodes* are defined as k-mers having in-degree or out-degree greater than 1. *Terminal nodes* are k-mers having out-degree 1 and in-degree 0 or in-degree 1 and out-degree 0. Terminals and junctions are collectively referred to as *special nodes*. The *compacted de Bruijn graph* is obtained from a de Bruijn graph by merging all adjacent *non-branching nodes* (i.e., those having in-degree and out-degree of exactly 1). The string associated with merged adjacent nodes is the first k-mer, concatenated with the single character extensions of all following non-branching k-mers. Such merged non-branching paths are called *unitigs*.

Since a junction *v* having in-degree greater than 1 and out-degree 1 is identified with *v′* having out-degree greater than 1 and in-degree 1, we speak of junction directions relative to the reading direction of the junction’s k-mer. Therefore, a *forward junction* has out-degree greater than 1, and a *back junction* has in-degree greater than 1. We refer to outbound k-mers beginning paths in the direction having out-degree greater than 1 as *heads*, and the sole outbound k-mer in the opposite direction as the junction’s *tail*. It is possible that a junction may have no tail.

A Bloom filter *B* is a space-efficient probabilistic hash table enabling insertion and approximate membership query operations [19]. The filter consists of a bit array of size *m*, and an element *x* is inserted to *B* by applying *h* hash functions, *f*_0_, …, *f_h−1_* such that *∀_i∈_*^[0*,h−*1]^*f*_*i*_(*x*) *∈* [0*, m −* 1], and setting values of the filter to 1 at the positions returned. For a Bloom filter *B* and string *s*, by *s ∈ B* or the term ’s in B’ we refer to *B*[*s*] = 1, i.e., when the *h* hash functions used to load *B* are applied t o *s*, only 1 values are returned. Similarly, *s ∉ B* or ’s not in B’ means that at least one of the *h* hash functions of *B* returned 0 when applied to *s*. For any *s* that has been inserted to *B, B*[*s*] = 1 by definition (i.e., there are no false negatives). However, false positives are possible, with a probability that can tuned by adjusting *m* or *h* appropriately.

## 3 Methods

We developed an algorithm called Faucet for streaming de novo assembly graph construction. A bird’s eye view of its entire work-flow is provided in Figure 1. Below we detail individual steps.

**Fig. 1:**
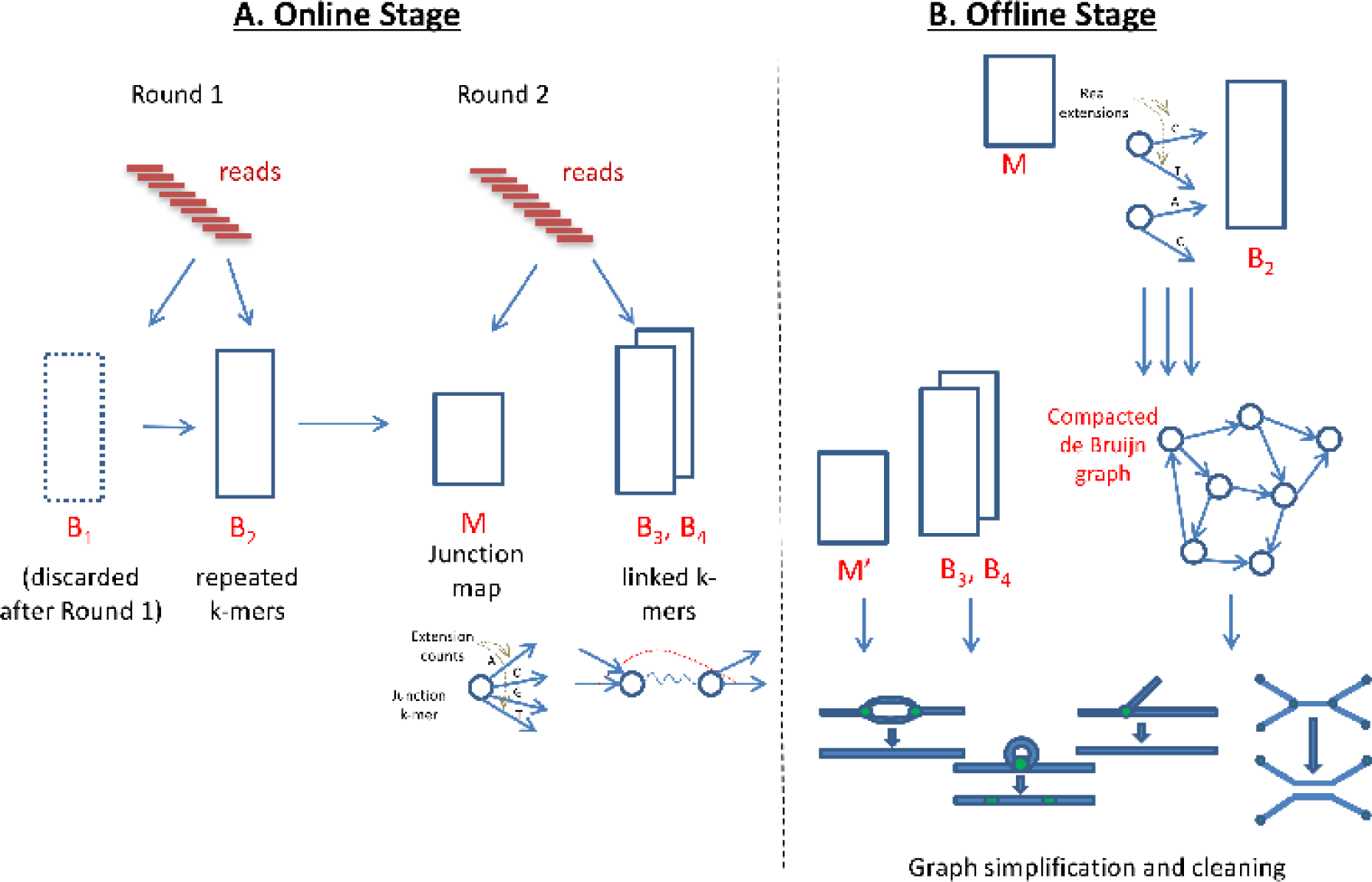
Faucet work-flow. A. The online stage involves a first round of processing all reads in order to load Bloom filters *B*_1_ and *B*_2_, and a second round in order to build the junction map M and load additional Bloom filters *B*_3_ and *B*_4_. *M* stores the set of all junctions and extension counts for each junction, while *B*_3_ and *B*_4_ capture connections between junction pairs. The two online rounds capture information from and perform processing on each read, and the processing performed always depends on the current state of data structures being loaded. B. The offline stage uses *B*_2_ and *M*, constructed during the online stage, in order to build the compacted de Bruijn graph by extending between special nodes using Bloom filter queries. ContigNodes (not shown) take the place of junctions and are stored in *M*′, allowing access (via stored pointers) to Contigs out of each junction, and coverage information. An additional vector of coverage values at fake or past junctions is also maintained for each Contig. Then, *B*_3_, *B*_4_, and this coverage information are used together to perform simplifications on and cleaning of the graph.

### Online Bloom Filter Loading

Faucet begins by loading two Bloom filters, *B*_1_ and *B*_2_, as it iterates through the reads, using the following procedure: all k-mers are inserted to *B*_1_, and only k-mers already in *B*_1_ (i.e., those for which all hash queries return 1 from *B*_1_) are inserted to *B*_2_. Namely, for each k-mer *s*, if *B*_1_[*s*] = 1 then we insert *s* into *B*_2_, otherwise we insert into *B*_1_. After iterating through all reads, *B*_1_ is discarded and only *B*_2_ is used for later stages. This procedure imposes a coverage threshold on the vast majority of k-mers so that primarily ’solid k-mers’ [20] observed at least twice are kept. This process is depicted in Round 1 of Figure 1A. We note that a small proportion of singleton or false positive k-mers may evade this filtration. No count information is associated with k-mers at this round.

### Online Graph Construction

*B*_2_, loaded at the first round, enables Faucet to query possible forward extensions of each k-mer. Faucet iterates through all reads a second time to collect information necessary for avoiding false positive extensions, building the compacted de Bruijn graph, and later, cleaning the graph. The second round consists of finding junctions and terminal k-mers, recording their true extension counts, and recording k-mer pairs (Round 2 of Figure 1A).

Faucet’s Online stage has one main routine - Algorithm 1 - that calls upon two subroutines - Algorithm 2 and Algorithm 3. First, junction k-mers and their start positions are derived from a call to Algorithm 2. To find junctions, Algorithm 2 makes all possible alternate extension queries (Line 3-Line 4) to *B*_2_ for each k-mer in the read sequence *r*. A junction k-mer *j* may have multiple extensions in *B*_2_ - either because there are multiple extensions of *j* in *G* that are all real (i.e., present on some read), or because there is at least one real extension in *G* and some others in *B*_2_ that are false positives. Accordingly, each k-mer possessing at least one extension that differs from the next base on the read is identified as a junction. Whenever one is found, its sequence along with its start position are recorded (Line 4), and the list of such tuples is returned. We note that each k-mer in the read is also queried for junctions in the reverse complement direction, but this is not shown in Algorithm 2.

Algorithm 1 then uses this set of junctions to perform accounting (Line 4-Line 7). All junctions are inserted into a hash map *M* that maps junction k-mers to vectors maintaining counts for each extension. For each junction of *r*, a count of 0 is initialized for each possible extension. These counters are only incremented based on extensions observed on reads - i.e., extensions due to Bloom filter outputs alone are not counted. As every real extension out of each junction must be observed on some read, and we scan the entire set of reads, an extension will have non-zero count only if it is real. This mechanism allows Faucet to maintain coverage counts for all real extensions out of junctions. In later stages, only extensions having non-zero counts will be visited, but counts are stored for real extensions of false junctions as well. These latter counts are used to sample coverage distributions on unitig sequences at more points than just their ends. Proportions of real junctions vs. the totals stored after accounting are described in the section ’Solid junction counts’ in the Appendix.

#### Algorithm 1 *scanReads*(*R, B*_2_)

~~~
**Input:** read set *R*, Bloom filter *B*_**2**_loaded from round 1, an empty Bloom filter *B*_**3**_
~~~

~~~
**Output:** 1. a junction Map *M* comprised of (*key, value*) pairs. Each *key* is a junction k-mer, and each *value ∈* N4 is a vector [*cA, cC, cG, cT* ] of counts representing the number of times each possible extension of *key* was observed in *R*; 2. *B*_**3**_ is loaded with linked k-mer pairs (i.e., specific 2k-mers - see text - are hashed in).
~~~

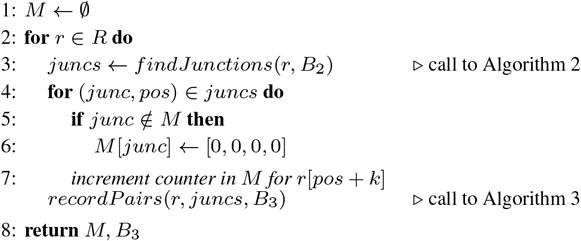

#### Algorithm 2 *findJunctions*(*r, B*_**2**_)

~~~
**Input:** read *r* and Bloom filter *B*_**2**_
~~~

~~~
**Output:** juncTuples, a list of tuples (*seq, p*), where *p* is the start position of junction k-mer *seq* in *r*, in order of appearance on *r*
~~~

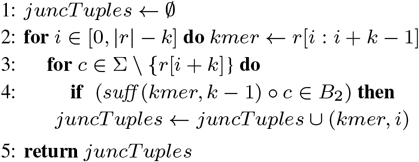

Following the accounting performed on observed junctions, Faucet records adjacencies between pairs of junctions using additional Bloom filters - *B*_3_ and *B*_4_. These adjacencies are needed for disentanglement - a cleaning step applied in Faucet’s offline stage. Disentanglement, depicted in Figure 2, is a means of repeat resolution. Its purpose is to split paths that have been merged due to the presence of a shared segment - the repeat - in both paths. In order to ’disentangle,’ or resolve the tangled region into its underlying latent paths, we seek to store sequences that flank opposite ends of the the repeat. Pairs of heads observed on reads provide a means of ’reading out’ such latent paths by indicating which heads co-occur on sequenced DNA fragments. The application of disentanglement is presented in the section ’Offline graph simplification and cleaning,’ while we now focus on the mechanism of pair collection and its rationale. To capture short and long range information separately, Bloom filter *B*_3_ holds head pairs on the same read, while *B*_4_ holds heads chosen such that each head is on a different mate of a paired-end read. Algorithm 3 is the process by which pairs are inserted into *B*_3_, and insertion into *B*_4_ is described in the Appendix.

**Fig. 2:**
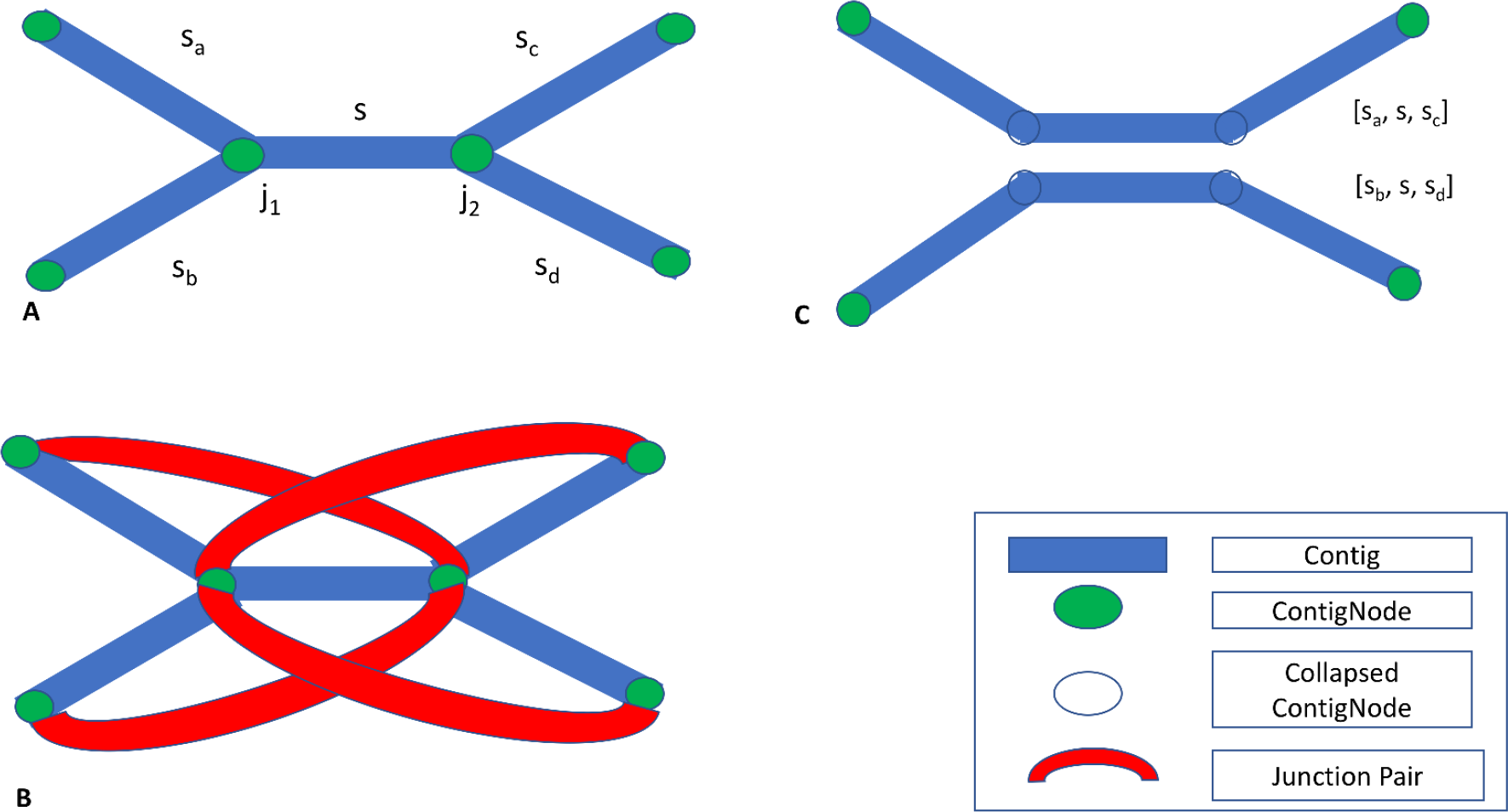
Disentanglement. A. A *tangle* characterized by two opposite facing junctions *j*_1_ and *j*_2_, each with out-degree 2. B. Junction pairs linking extensions on *s*_*a*_ with *s*_*c*_ and *s*_*b*_ with *s*_*d*_. Since no pairs link extensions on *s*_*a*_ with *s*_*d*_ or *s*_*b*_ with *s*_*c*_, only one orientation is supported. C. the result of disentanglement: paths [*s*_*a*_*, s, s*_*c*_ ] and [*s*_*b*_*, s, s*_*d*_ ] are each merged into individual sequences, and junctions *j*_1_ and *j*_2_ are removed from *M*.

In Algorithm 3, we aim to pair heads that are maximally informative. Informative pairs are those that allow us to ’read out’ pairs of unitigs that belong to the same latent path. We specifically choose to insert heads because during the offline stage when disentanglement takes place, adjacencies between each unitig starting at an edge to a head and the unitig starting at the edge from the junction to its tail of are known and accessible via pointers to their sequences. Therefore, extension pairs capturing information of direct adjacencies provide no new information. The closest indirect adjacency that may be informative when captured from a read is that between two junctions that either face in the same direction, or when the first faces back and the second faces forward, as shown in Figure 3 A. Thus, when there are only two junctions on a read, their pair of heads is inserted as long as the two junctions are not facing each other. When there are at least three junctions on a read, every other junction out of every consecutive triplet is paired, as shown for a single triplet in Figure 3B. This figure demonstrates that selecting every other head is preferable to selecting consecutive heads out of a triplet. This type of insertion is executed in Line 1-Line 5 of Algorithm 3 and ensures all unitigs flanking some triplet are potentially inferable. For reads having more than three junctions, applying the triplet rule for every consecutive window of size 3 similarly allows for all unitigs on the read to be included in some hashed pair.

**Fig. 3:**
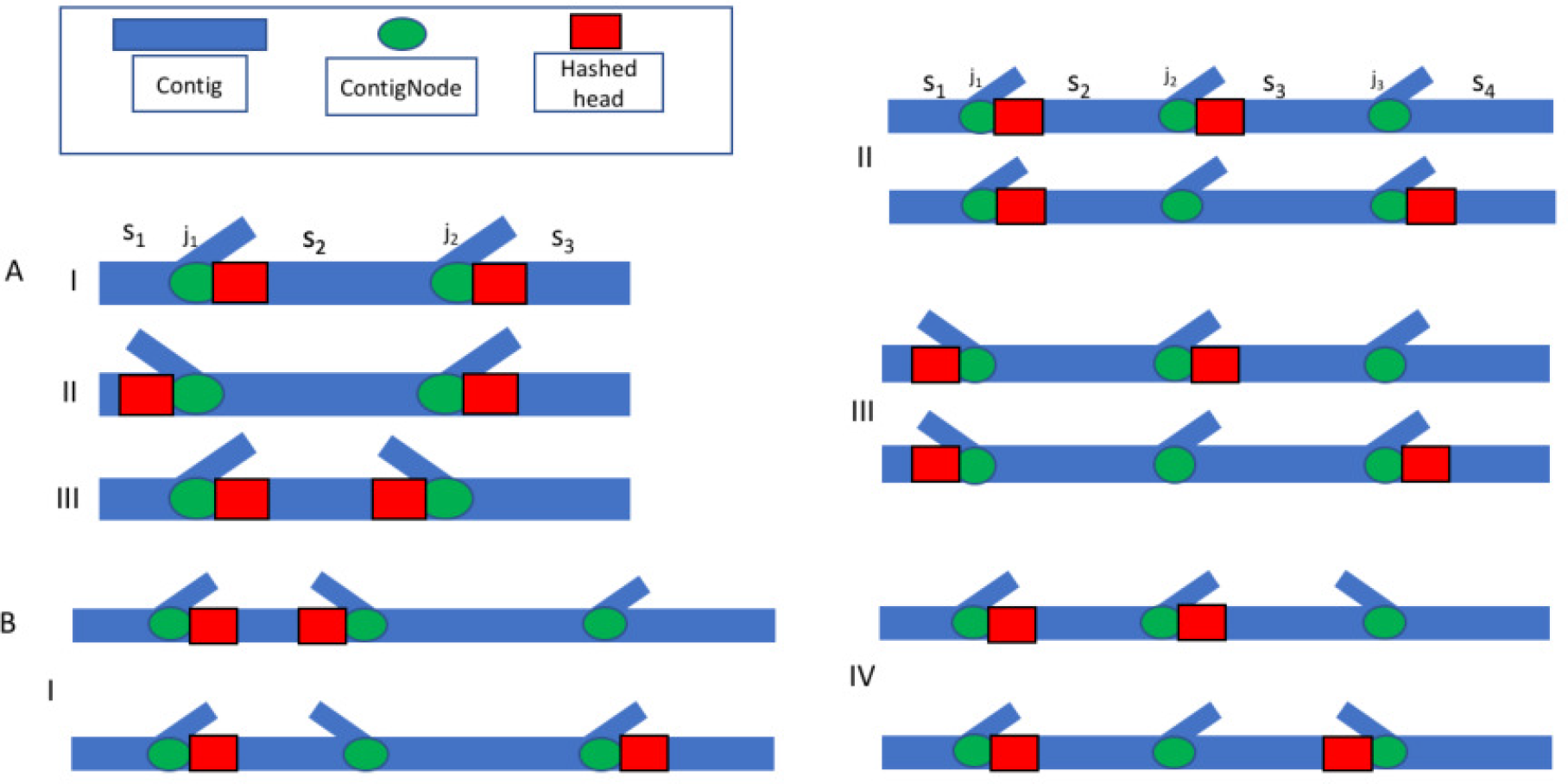
Rationale for *B*_3_ insertions. Pairs of junction heads (indicated by red rectangles adjacent to green junctions) observed on reads are inserted when they provide additional information to infer a path on the graph. A. Two junctions observed on a read. In cases I and II, it is beneficial to insert the pairs of heads into *B*_3_, as in both cases each individual head allows inference of a different Contig. In case III, inserting the pair between two junctions facing each other is not beneficial because both heads lie on opposite ends of the same Contig. In all three cases, the full path can be inferred. B. Four possible arrangements of three consecutive junctions on a read. There are four more that are symmetrical reflections of those presented that are not shown. In each case, we compare the Contigs covered (i.e., either included by some head or inferable as a junction’s back) when heads out of consecutive (top) and non-consecutive (bottom) junctions are chosen, assuming only one pair is inserted. Note that in cases I-III, Contig *s*_4_ is not covered by any paired head or tail when inserting consecutive heads, while in case IV, all Contigs are covered by either the paired heads or some tail. Thus, in the first three cases it will not be possible to determine which head of *j*_3_ occurred with an extension of *j*_1_ or *j*_2_ on some read unless this information is provided by some other hashed pair. In contrast, when non-consecutive heads are paired, every Contig is covered by either one of the inserted heads or a tail.

### Offline Graph Simplification and Cleaning

Given *B*_2_, *B*_3_, *B*_4_ and *M* resulting from the online stage, the compacted de Bruijn graph is generated by traversing each forward extension out of every special k-mer, as well as traversing backwards in the reverse complement direction when the node has not been reached before by a traversal starting from another node. This is done by querying *B*_2_ for extensions and continuing until the next special node is reached. During each such traversal from special node *u* to special node *v*, a unitig sequence *s*_*uv*_ is constructed. *s*_*uv*_ is initialized to the sequence of *u*, and a base is added at each extension until *v* is reached.

#### Algorithm 3 *recordPairs*(*r, juncs, B*_3_)

~~~
**Input:** read *r, juncs* - a list of pairs (*j, p*), where *p* is the start position of junction *j* in *r*, and Bloom filter *B*_3_. We also make use of a subroutine *getOutExt*(*j*_*i*_*, p*_*i*_*, r*) that for a junction *j*_*i*_ returns *pref* (*j*_*i*_*, k −* 1) ∘ *r*[*p*_*i*_ *− k*] if *j*_*i*_ is a back junction, and *suff* (*j*_*i*_*, k −* 1) ∘ *r*[*p*_*i*_ + *k*] otherwise.
~~~

~~~
**Output:** Bloom filter *B*_3_, loaded with select *linked k-mer pairs*
~~~

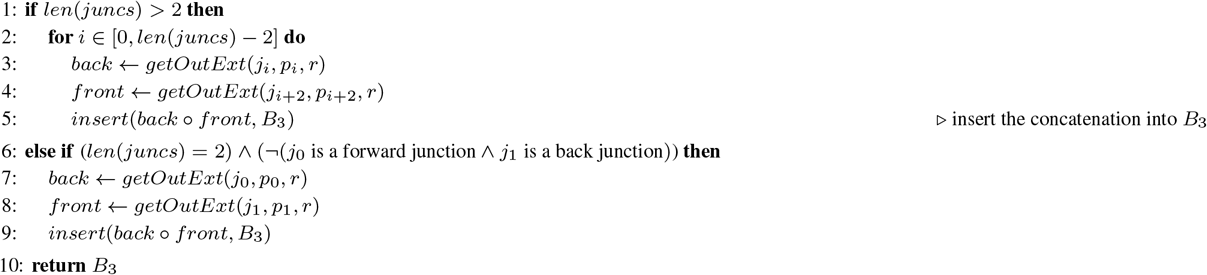

New data structures are constructed in the course of traversals in order to aid later queries and updates. A *ContigNode* structure is used to represent a junction that points to *Contigs*. ContigNodes are structures possessing a pointer to a Contig at each forward extension, as well as one backwards pointer. This backwards pointer connects the junction to the sequence beginning with the reverse complement of the junction’s k-mer. Contigs initially store unitig sequences, but these may later be concatenated or duplicated. They also point to one ContigNode at each end. To efficiently query Contigs and ContigNodes, a new hashmap *M*′ is constructed having junction k-mers as keys, and ContigNodes that represent those junctions as values. Isolated contigs formed by unitigs that extend between terminal nodes are stored in a separate set data structure.

Once the raw graph is obtained, cleaning steps commence, incorporating tip removal, chimera removal, collapsing of bulges, and disentanglement. Coverage information and paired-junction links are crucial to these steps. Briefly, tip removal involves deletion of Contigs shorter than the input read length that lead to a terminal node. Chimera and bulge removal steps involve heuristics designed to remove low coverage Contigs when a more credible alternative (higher coverage, or involved in more sub-paths) is identified. These first three steps proceed as described in [21], thus we omit their full description here.

Disentanglement relies on paired junction links inserted into *B*_3_ and *B*_4_. We iterate through the set of ContigNodes to look for ’tangles’ - pairs of opposite-facing junctions joined by a repeat sequence - as shown in Figure 2. Tangles are characterized by tuples (*j*_1_*, j*_2_*, s*) where *j*_1_ is a back junction, *j*_2_ is a forward junction (or vice-versa), and there is a common Contig *s* pointed to by the back pointers of both *j*_1_ and *j*_2_. Junctions *j*_1_ and *j*_2_ each have at least two outward extensions. We restrict cleaning to tangles having exactly two extensions at each end. Let *s*_*a*_ and *s_b_* be the Contigs starting at heads of *j*_1_, and *s*_*c*_ and *sd* be the Contigs starting at heads of *j*_2_. By disentangling, we seek to pair extensions at each side of *s* to form two paths. The possible outputs are paths [*s*_*a*_*, s, s*_*c*_] together with [*s_b_, s, s_d_*] or [*s*_*a*_*, s, s_d_*] together with [*s_b_, s, s*_*c*_].

Thus, each such pair straddling the tangle -e.g., having one head on *s*_*a*_ and the other on *s*_*c*_ - lends some support to the hypothesis that the correct split is that which pairs the two. To decide between the two possible split orientations, we count the number of pairs supporting each by querying *B*_3_ or *B*_4_ for all possible junction pairings that are separated by a characteristic length associated with the pairs inserted to each. For example, *B*_3_ stores heads out of non-consecutive junction pairs on the same read. Therefore, for each junction on *s*_*a*_ we count each pairing accepted by *B*_3_ with a junction on *s*_*c*_ that is at most one read length away. Specifically for *B*_3_, we also know that inserted pairs are always one or two junctions away from the starting junction, based on the scheme presented in Figure 3. To decide when a tangle should be split, we apply XOR logic to arrive at a decision: if the count of pairs supporting both paths in one orientation is greater than 0, and the count of both paths in the other orientation is 0, we disentangle according to the first, as shown in Figure 2. Similar yet more involved reasoning is used for junction links in *B*_4_, using the insert size between read pairs (see Appendix). Once we arrive at a decision, we add a new sequence to the set of Contigs that is the concatenation of the sequences involved in the original paths. We note one of the consequences of this simplification step is that the graph no longer represents a de Bruijn graph, in that each k-mer is no longer guaranteed to appear at most once in the graph. Furthermore, the XOR case presented is the most frequently applied form of disentanglement out of a few alternatives. We discuss these alternatives in the Appendix.

### Optimizations and Technical Details

Here we discuss some details omitted from the above descriptions for the sake of completeness. Based on the description of Algorithm 1 and Algorithm 2, it is possible that false positive extensions out of terminal nodes will ensue. This is possible because the mechanism described for removing false positive junctions can differentiate between one or multiple extensions existing in *G* for a given node, but can not differentiate between one or none. This may lead to assembly errors at sink nodes.

To overcome such effects, we store distances between junctions seen on the same read with the distance recorded being assigned to the extension of each junction observed on the read. When an outermost junction on a read has not been previously linked to another junction, we record its distance from the nearest read end - this solves the problem mentioned previously as long as paths to sinks are shorter than read length. To obtain accurate measurements of distances on longer non-branching paths, we also introduce artificial ’dummy’ junctions whenever a pre-defined length threshold is surpassed. In effect, this means that reads with no real junctions are assigned dummy junctions.

Once distances and dummy junctions are introduced, an additional benefit is gained: the speed of the read-scan can be improved by skipping between junctions that have been seen before. Once distances are known, if we see a particular extension out of a junction, and then a sequence of length ℓ without any junctions, then, wherever else we see that junction and extension, it must be followed by the exact same ℓ next bases. Otherwise, there would be a junction earlier. So we store *f* when we see it, and skip subsequent occurrences.

Finally, we note that Faucet can benefit from precise Bloom filter sizing. When a good estimate of dataset parameters is known, the algorithm can do the 2-pass process above. Otherwise, to determine the numbers of distinct k-mers and the number of singletons in the dataset in a streaming manner, we have used the tool ntCard [15]. This requires an additional pass over the reads (for a total of three passes). The added pass does not increase RAM or disk use. In fact, in tests on locally stored data, we found it only adds negligible time.

## 4. Results

### Assembling While Downloading

As a demonstration of streaming assembly, we ran Faucet on publicly available human data, SRR034939, used for benchmarking in [6]. To assess resource use at different data volumes, we ran Faucet on 10, 20, and 37 paired-end files out of 37 total. Streaming was enabled using standard Linux command line tools: wget was used for commencing a download from a supplied URL, and streamed reading from the compressed data was enabled by the bzip2 utility. Downloads were initiated separately for each run. The streaming results are shown in Table 1.

**Table 1.**

| No. of files | Time (hrs) | RAM (GB) | Disk (GB) | Data size (GB) | Comp. ratio |
| --- | --- | --- | --- | --- | --- |
| 10 | 26.3 | 48.3 | 19.0 | 29.6 | 0.64 |
| 20 | 47.7 | 84.3 | 34.3 | 59.2 | 0.58 |
| 37 | 98.2 | 144.7 | 50.0 | 108.4 | 0.46 |

We emphasize that Faucet required less space than the size of the input data in order to assemble it, while most assemblers generate files during the course of their processing that are larger than the input data. Also, the ratio of input data to disk used by Faucet decreased as data volume increased, reflecting the tendency of sequences to be seen repeatedly with high coverage. We also note that Faucet’s outputs effectively create a lossy compression of the read data, in that the choice of k value inherently creates some ambiguity for read substrings larger than k. This compression format is also queryable, in that given a k-mer in the graph, its extensions can be found: indeed, this is the basis of Faucet’s graph construction and cleaning.

### Disentanglement Assessment

To gauge the benefits of disentanglement on assembly quality, we compared Faucet’s outputs with and without each of short- and long-range pairing information, provided by Bloom filters *B*_3_ and *B*_4_, on SYN 64 - a synthetic metagenome produced to provide a dataset for which the ground truth is known comprised of 64 species (data set sizes and additional characteristics are provided in the Appendix). The results of this assessment are presented in Table 2. We measured assembly contiguity by the NGA50 measure. NG50 is defined as “the contig length such that using equal or longer length contigs produces x% of the length of the reference genome, rather than x% of the assembly length” in [22]. NGA50 is an adjustment of the NG50 measure designed to penalize contigs composed of misassembled parts by breaking contigs into aligned blocks after alignment to the reference. We found that disentanglement more than doubled contiguity measured by mean NGA50 values, with greater gains as more kinds of disentanglement were enabled. This was also reflected by corresponding gains in the genome fractions, and in the number of species for which at least 50% of the genome was aligned to, allowing NGA50 scores to be reported. More applications of disentanglement also increased the number of misassemblies reported and the duplication ratio, however two thirds of the maximum misassembly count is already seen without any disentanglement applied.

**Table 2.**

| | No disent. | $B_3$ only | $B_4$ only | both $B_3, B_4$ |
| --- | --- | --- | --- | --- |
| Genome fraction (%) | 76.4 | 79.9 | 80.3 | 82.3 |
| Dup. ratio | 1.00 | 1.01 | 1.02 | 1.02 |
| Mean NGA50 | 13048 | 21703 | 26356 | 29066 |
| Misassemblies | 388 | 480 | 521 | 572 |
| Species reported | 54 | 56 | 56 | 56 |

### Tools Comparison

We sought to assess Faucet’s effectiveness in assembling metagenomes, and its resource efficiency. For the former, we compared Faucet to MetaSPAdes [23] and Megahit [24], state of the art metagenome assemblers in terms of contiguity and accuracy that require substantial resources. To address resource efficiency, we also compared Faucet to two leading resource efficient assemblers, Minia 3 (Beta) [6] and LightAssembler [17]. We note these last two were not designed as metagenome assemblers, but they perform operations similar to what Faucet does - both in the course of their graph construction steps, and in their cleaning steps. They differ from Faucet in that neither is capable of disentanglement, as they do not utilize paired-end information, but counter this advantage with more sophisticated traversal schemes. All tools were run on two metagenome data sets - SYN64 and HMP - a female tongue dorsum sample sequenced as part of the Human Microbiome Project. Both datasets were used for testing in [23]. To achieve a fair comparison, runs were performed with a single thread on the same machine, as Faucet does not currently support multi-threaded execution. Full details of the comparison, including versions, parameters, and data accessions, are presented in the supplement.

Table 3 presents the full results for the tools comparison. There was a strong advantage to Megahit and MetaSPAdes over the three lightweight assemblers (Minia, LightAssembler, and Faucet) in terms of contiguity achieved (shown by NGA50 statistics), but this came at a large cost in terms of memory, disk space, and time, particularly in the case of MetaSPAdes. Among the lightweight assemblers, Minia used by far the most disk space, and differences in other resource measures were less pronounced. Among these three, Faucet had a large advantage in NGA50 statistics relative to the other two. This is highlighted by the trend of Table 3, and shown by its 14-110% advantage in the mean of NGA50 relative to Minia, and 2-11 *fold* advantage relative to LightAssembler.

**Table 3.** Tool comparison on two metagenomes. Top values in each cell are for SYN 64 data, and bottom values are for HMP. Duplication ratio is the ratio between the total aligned length to the combined length of all references aligned to. The mean and median NGA50 values are calculated on based on species sufficiently covered by all assemblers to yield an NGA50 value (i.e., 50% of the genome is covered). Species reported are those for which an NGA50 value is reported. In the HMP data, only 2 species were reported for all, making the mean and median NGA50 values equal. Disk and memory use are those reported by the Linux time utility, and Disk use is the total amount written to disk during the course of a run.

|  | Metaspades | Megahit | LightAssembler | Minia | Faucet |
| --- | --- | --- | --- | --- | --- |
| Genome fraction (%) | 89.1 | 90.1 | 75.6 | 76.5 | 82.3 |
| Dup. ratio | 1.02 | 1.02 | 1.01 | 1.00 | 1.02 |
| Mean NGA50 (kb) | 167 | 99.0 | 2.60 | 14.6 | 30.7 |
| Median NGA50 (kb) | 71.1 | 57.6 | 2.30 | 10.5 | 23.7 |
| Misassemblies | 785 | 949 | 314 | 395 | 572 |
| Species reported | 59 | 61 | 55 | 52 | 56 |
| Time (hrs) | 41.2 | 10.9 | 1.63 | 0.97 | 2.61 |
| Memory (GB) | 26 | 9.1 | 2.7 | 4.8 | 6.0 |
| Disk (GB) | 43.1 | 14.3 | 28.2 | 1.84 | 1.59 |
| Genome fraction (%) | 46.9 | 48.6 | 23.4 | 27.8 | 27.9 |
| Dup. ratio | 1.05 | 1.12 | 1.02 | 1.01 | 1.05 |
| Mean NGA50 (kb) | 28.3 | 36.8 | 3.18 | 6.25 | 7.12 |
| Median NGA50 (kb) | 28.3 | 36.8 | 3.18 | 6.25 | 7.12 |
| Misassemblies | 504 | 602 | 100 | 184 | 202 |
| Species reported | 12 | 12 | 5 | 3 | 6 |
| Time (hrs) | 30.5 | 13.0 | 3.35 | 0.99 | 2.30 |
| Memory (GB) | 14 | 8.3 | 3.4 | 3.7 | 7.3 |
| Disk (GB) | 53.2 | 11.5 | 23.5 | 1.30 | 1.61 |

**Table 4.**

|  | HMP | SYN 64 |
| --- | --- | --- |
| Total junctions (M) | 7.11 | 9.23 |
| Real junctions (M) | 4.55 | 1.34 |
| genome fraction (%) | 27.9 | 82.3 |
| N50 | 2290 | 16707 |

## 5. Discussion

Streaming de novo assembly presents an opportunity to significantly ease some of the burdens introduced by the recent deluge of second generation sequencing data. We posit the main applications of streaming assembly will be de novo assembly of very large individual datasets (e.g., metagenomes from highly diverse environments) and re-assembly of pangenomes derived from many samples. In both cases, very large volumes of data must be digested in order to address the relevant biological questions behind these assays. Therefore, streaming graph assembly presents an attractive alternative to data compression: instead of attempting to reduce the size of data, the aim is to keep locally only relevant information in a manner that is queryable and that allows for future re-analysis.

Here, we have demonstrated a mechanism for performing streaming graph assembly and described some of its characteristics. First, we showed that assembly can be achieved without ever storing raw reads locally. By assembling the graph, an intermediate by-product of many assemblers, we show this technique is generally applicable. By refining the graph and showing better assembly contiguity than competing resource efficient tools on metagenome assembly, we showed this method can also be applied in the setting when sensitive recovery of rare sequences is crucial.

In future work, we aim to expand the capabilities of Faucet in a number of ways. Multi-threaded processing will reduce run times and make the tool more applicable to large data volumes. We believe further refinements of cleaning and contig generation can be achieved by adopting a statistical approach to making assembly decisions. In addition, beyond graph cleaning, we aim to apply Faucet’s data structures to path generation, as done with paired end reads in [25, 26, 27]. Both have the potential to greatly improve contiguity and accuracy.

Beyond this, the present work raises several remaining challenges pertaining to what one may expect of streaming assembly. For instance, it is immediately appealing to ask if streaming assembly can be achieved with a just a single pass on the reads, and if so, what inherent limitations exist. In [12], a simple solution is proposed wherein the first 1M reads are processed to provide a succinct summary for the rest, but such an approach is more suited to high coverage or low entropy data, and thus unlikely to perform well on diverse metagenomes or when rare events are of particular interest. Another issue raised by the performance comparison herein is that of capturing the added value that iterative (multi-k value) graph generation provides. We have given a partial solution by capturing subsets of junction pairs within each read, and between mates of paired-end reads. Although it is possible to iteratively refine the graph with more passes on the reads, each time for the collection of k-mers at different lengths, this becomes unwieldy with large data volumes. Identifying the contexts for which such information would be useful in the graph and indexing the reads to allow for querying of such contexts may provide more efficient means of extracting such information.

## 6.Acknowledgments

This work was supported in part by the Israel Science Foundation as part of the ISF-NSFC joint program to RS. RS was supported in part by the Raymond and Beverley Chair in Bioinformatics at Tel Aviv University. EH was supported in part by the United States-Israel Binational Science Foundation (Grant 2012304) and EH and RR were supported in part by the Israel Science Foundation (Grant 1425/13). RR was supported in part by a fellowship from the Edmond J. Safra Center for Bioinformatics at Tel Aviv University, an IBM PhD fellowship, and by the Center for Absorption in Science, the Israel Ministry of Immigrant Absorption. EH is a Faculty Fellow of the Edmond J. Safra Center for Bioinformatics at Tel Aviv University. GG was supported by the MISTI MIT-Israel program at MIT and Tel Aviv University.

## Appendix

### Sizing Bloom Filters

We used the tool ntCard to estimate the cardinality *F*_0_ of the set of k-mers and the number of singletons *f*_1_. These counts are used for optimizing both runtime and memory use by allowing us to minimize the size of Bloom filters *m* and the number of hash functions *h* used for both *B*_1_ and *B*_2_, the largest filters used. *B*_1_ and *B*_2_ share these parameters (and the same set of hash functions) to allow insertions of each *s* into *B*_2_ for which *B*_1_(*s*) = 1 without recalculating *h* hash values. We use the fact that elements inserted into *B*_2_ are either non-singletons or false positives due to *B*_1_. Thus, the expected number of elements *n*_2_ in *B*_2_, is bound by their sum, i.e.,

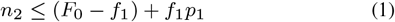
where *p*_1_ is the false positive rate of *B*_1_. We note that since *p* is the *effective* false positive rate after all elements are inserted into *B*_1_, this bound holds strictly and may be overly pessimistic regarding the number of false positives inserted into *B*_2_, however it provides a simple means of setting parameters. To do so, we first recall that *B*_1_ is discarded after loading, while *B*_2_ is maintained and thus its false positive rate *p*_2_ is the rate that affects all downstream queries. A default false positive rate of of *p*_2_ = 0.01 is used to work backwards to derive a higher rate *p*_1_, and Bloom filter parameters for both filters were set based on this derived value, using knowledge of *F*_0_ and *f*_1_. To derive *p*_1_, we paired the expressions for the expected false positive rates with the expression for the optimal number of hash functions for a given false positive rate [28]:

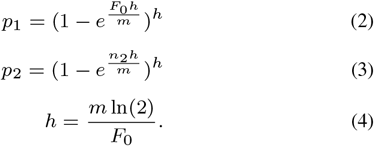
By plugging the value of *h* from equation 4 into equation 2, we arrive at 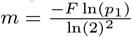. Combining this and the above expressions, we arrive at

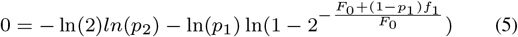
for which root-finding methods can be applied to finally extract *p*_1_, the sole remaining unknown.

Currently, we have not yet found similar means of optimizing the sizes of filters *B*_3_ and *B*_4_, as it is unclear how to estimate the number of elements that will be inserted into them in advance. We therefore define their sizes based on empirical observations. For diverse metagenomes, where the number of singletons *f*_1_ may be very close to the cardinality *F*_0_, we expect there to be few junctions, as a junction k-mer must by definition occur at least twice in the data. Based on this observation, we set the expected number of elements in both *B*_3_ and *B*_4_ to be 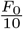 and found that this bound was not exceeded on tested datasets. For higher coverage data, where a significantly larger proportion of junctions is expected relative to *F*_0_, we set the size of both filters to be 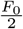.

### Solid Junction Counts

Total junction counts listed in the table below include real junctions, those due to false positives, and dummy junctions inside long linear stretches. We posit that the SYN 64 data set included many more fake (false positive and dummy) junctions as a result of having a much larger proportion of linear stretches, as reflected in the much larger genome fraction and N50 size (relative to HMP) output by Faucet.

### Inserting into *B*_4_

When inserting into *B*_4_, both the distance and relative orientation between paired-end mates is unknown. Therefore, a tiling scheme such as that seen in Figure 3 cannot be applied. Instead, we seek to ensure that in most cases when querying approximately one insert size away from a given junction *u*, there will be another junction *v* such that an extension of *u* will be paired with an extension of *v* in *B*_4_. To achieve this end, and to avoid long run times due to pair insertions, we apply the following logic: for each junction *u* on the first mate, we only insert extensions of a new pair (*u, v*) if *u* has no pair in *B*_4_. When a new pair must be inserted, *v* is chosen to be the first junction found on the second mate. During the insertion process, this logic allows us to break the querying process whenever one previously inserted pair is encountered, and lets us avoid inserting too many pairs into *B*_4_, and thus risking increasing *B*_4_’s effective false positive rate.

### Additional Disentanglement

Other forms of disentanglement include resolution of loops and disentanglement by coverage. Loops are encountered when, e.g., *s*_*a*_ and *s*_*c*_ in Figure 2 are the same unitig, and disentanglement requires unwinding the loop and duplicating the *s*’s sequence to arrive at the walk [*s_b_, s, s*_*c*_*, s, s_d_*]. Disentanglement by coverage is allowed only when *s* is deemed too long for there to be support by junction pairs flanking opposite ends of *s*, and is applied when the coverage distributions of Contig pairs supporting a certain orientation (e.g., *s*_*a*_ paired with *s*_*c*_ and *sb* with *sd* for the case presented in Figure 2) is significantly similar, as determined by Two One Sided Tests [29] for each pair. To smooth coverage levels when this test is applied, coverage values are updated each time a cleaning step such as bulge removal is applied. For example, if a bubble includes one low coverage Contig *s*_1_ and one high coverage Contig *s*_2_, as extensions flanked by the same ContigNodes *jL* and *jR*, and *s*_2_’s coverage is sufficiently higher than *s*_1_’s, Contig *s*_1_ will be removed, and its average coverage will be assigned to all (expired or fake) junctions on Contig *s*_2_.

### Tools Comparison Details

Tools and flags:

Faucet was run with k= 31

MetaSPAdes 3.9.0, default parameters

Megahit 1.1.1, default parameters

Minia 3 Beta, git commit 4b0a83a, k = 31

LightAssembler, no version information available, downloaded 1/17 from GitHub k = 31

MetaQUAST, 4.4.0, –fragmented flag

Data Sets:

SYN 64 (SRA accession SRX200676), 109M 100 bp paired end mates, I.S. 206

HMP (SRX024329), 149.6 M 100 bp paired end mates, I.S. 213

